# Impact of Voluntary Wheel Running on Gene Expression of Histone 3 Lysine 27-Modifying Enzymes in Mouse Skeletal Muscle

**DOI:** 10.1101/2025.11.07.687293

**Authors:** Daniel Gamu, Makenna Cameron, Yasmeen Jalil, Imen Zouaoui, Jada Sangha, Jonathan Kim

**Affiliations:** School of Kinesiology, University of British Columbia; Department of Medical Genetics, University of British Columbia

**Keywords:** exercise, skeletal muscle, histone acetylation/methylation, muscle adaptation

## Abstract

Skeletal muscle is highly plastic and capable of remodelling its contractile and metabolic properties depending on physical demands. Such remodelling requires modification of chromatin structure to support transcriptional activation and suppression of gene programs. Chromatin dynamics depend, in part, on the acetylation and methylation of histone 3 lysine 27 (H3K27), which is controlled by several H3K27-specific acetyltransferases, deacetylases, methyltransferases and demethylases. Several histone post-translational modifications in muscle have been shown to be modulated by exercise. Here, we sought to examine whether major H3K27 regulators are altered by endurance training. Male and female C57BL/6J mice were provided with voluntary running wheels for 6 weeks and compared to sex-matched sedentary controls with locked running wheels. Gene expression of various slow/oxidative and fast/glycolytic skeletal muscles was then measured. We found voluntary running induced modest changes in H3K27 acetyltransferases, along with several components of the polycomb-repressive complex 2 in a muscle- and sex-specific manner. Our findings indicate that the capacity for both acetylation and methylation of H3K27 is modulated by chronic endurance exercise, suggesting that chromatin dynamics are a mechanistic component of exercise-induced muscle remodelling.

## Introduction

Skeletal muscles comprise nearly half of adult body mass;^1^ as such, its functioning is vital to whole-body metabolism and overall health. At the molecular level, physical activity improves muscle functioning by remodelling a breadth of structural and metabolic programs, ultimately reducing disease risk. While much progress has been made in defining the intracellular signalling cascades governing muscle responsiveness to exercise, understanding precisely how exercise is decoded into an appropriate cellular adaptation remains incomplete. Such cellular remodelling requires modulation of gene programs by transcriptional activators and suppressors; however, transcription factor binding requires physical accessibility to the DNA within chromatin. To this end, much less is known about how the modification of chromatin structure controls skeletal muscle plasticity.

Within chromatin, DNA is tightly wrapped around histone proteins, which assist in nuclear packaging. Physical accessibility of transcriptional machinery to DNA, and thereby transcription, is regulated in part by epigenetic factors, including DNA methylation and post-translational modifications (PTMs) to histones. Histone (H) octamers are made of two copies each of H2A, H2B, H3 and H4 proteins. Among their numerous amino acid residues, histone 3 lysine 27 (H3K27) is particularly unique because it is subject to two mutually-exclusive PTMs with opposing effects on gene transcription.^2^ Acetylation (H3K27ac) facilitates chromatin “relaxation” in-part by neutralizing positively-charged N-terminal lysine residues to destabilize DNA-histone and histone-histone interactions, ultimately enabling binding of transcriptional machinery and transcriptional activation.^3,4^ Conversely, addition of up to 3 methyl groups (H3K27me1-3) results in tighter chromatin packaging and therefore, transcriptional silencing.^5^ The addition and removal of acetyl/methyl moieties is a complex function of several critical histone acetyl/methyltransferases and histone deacetyl/demethylases, respectively.

While histone PTMs have been well-described in the context of establishing cellular identity during development,^2^ their ongoing roles following organogenesis is underappreciated and unclear, particularly within highly adaptable tissues like skeletal muscle. Skeletal muscle can remodel various aspects of its biology in a stimulus-dependent manner. For example, endurance exercise training increases mitochondrial biogenesis and function, thereby enhancing the capacity for both muscle and whole-body lipid oxidation.^6^ Furthermore, endurance training can alter the expression of various contractile proteins, including increasing the proportion of “slow” vs “fast” contractile isoforms and ultimately muscle fibre-types.^6^ This enhancement in slow/oxidative programming contributes to the functional improvements in exercise performance and fatigue resistance with endurance training. While a number of signalling cascades and transcriptional master-regulators help co-ordinate muscle adaptations, mounting evidence indicates that histone PTMs, and thereby chromatin dynamics, play a central role.^7^

Genome-wide distribution of H3K27 PTMs differ among muscle fiber-types;^8,9^ however, they are not simply static across organismal lifespan. For example, histone acetylation/methylation is impacted by just a single exercise bout in both human and rodent muscle.^10–13^ Moreover, chronic endurance training can markedly alter genome-wide and gene-specific enrichment of H3K27ac and H3K27me1-3,^8,14,15^ likely supporting activation and repression of exercise-responsive gene programs. Importantly, endurance training can alter histone octamer composition and modulate expression of histone methylating enzymes.^16,17^ These findings suggest the capacity to change chromatin structure may be a critical regulator of muscle plasticity in response to exercise. Although factors determining acetylation and methylation of H3K27 are complex, whether or not the capacity to lay down or remove H3K27 PTMs is modulated by exercise training has yet to be examined. Here, we sought to examine if expression of H3K27ac/me1-3 regulatory enzymes fluctuate with endurance training adaptations across various mouse skeletal muscles.

## Materials and Methods

### Animals

Male and female C57BL/6J (10-16 week old) mice were used for all studies. Mice were housed at room temperature (∼22°C) under a 12hr-light/12hr-dark cycle in the animal facility at BC Children’s Hospital Research Institute (BCCHRI) and provided *ad libitum* access to water and standard rodent chow (Teklad-2918). All experiments were approved by the University of British Columbia Animal Care Committee in accordance with the Canadian Council on Animal Care guidelines.

### Voluntary Wheel Running (VWR)

All mice were group housed following weaning. Prior study commencement, littermates were randomly assigned to voluntary running wheels (VWR; male n = 8, female n = 7) or remained as sedentary controls (SED; male n = 8, female n = 6). Both VWR and SED mice were singly-housed in shoe-box style cages (height x width x length: 16 x 24 x 44 cm) containing food and water (as above), bedding material, a shelter, and an angled running wheel (Med Associates Inc; Fast-Trac^TM^ ENV-047 low-profile) connected to a wireless device hub (Med Associates Inc; DIG-807). SED controls had their wheels locked for the entire duration of the experiment. Acquisition of running wheel data was collected using Wheel Manager Software (Med Associates Inc; SOF-860). Running volume (km) was continuously monitored for 6 consecutive weeks, with physical activity data expressed as average daily running distance travelled within weeks 1 to 6 (i.e. km/day). Additionally, body mass (g) was measured at the beginning of each week. Week 4 activity and week 2 body mass data were missing for female mice. Following the experiment, all running wheels were locked for 48 hrs prior to tissue collection.

### Body Composition and Tissue Collection

Prior to (week 0) and immediately after (week 6) wheel running, body composition (i.e. lean and fat mass) was measured in non-anesthetized mice by quantitative magnetic resonance (EchoMRI-100H, Echo Medical Systems) as we have previously done.^18^ Forty-eight hours after wheel-lock, mice were anesthetized with an isoflurane/O_2_ mixture and euthanized by cervical dislocation. Visceral (gonadal and retroperitoneal) and subcutaneous (inguinal) adipose depots were removed and weighed; inguinal depots were collected along the entire inguinal crease, after which lymph nodes were removed. All skeletal muscles were cleared of visible blood and connective tissue prior to weighing and snap freezing in liquid nitrogen (LN_2_). Soleus and extensor digitorum longus muscles were excised from proximal to distal tendon. Red and white sections of the gastrocnemius and quadricep were partitioned by excising these visually distinct portions from the muscle belly, weighed and snap frozen in LN_2_. All skeletal muscles were then stored at −80°C until analysis.

### qPCR

Muscle samples were homogenized in TRIzol using a bead mill (Next Advance; Bullet Blender) and 0.5 mm zirconium oxide beads. Total RNA was column purified (RNeasy Mini Kit, Qiagen) according to manufacturer instructions, after which 0.5 µg of total RNA was reverse-transcribed (Invitrogen; SuperScript™ III, Cat. No 18080-051) using oligo(dT)_20_ primers. Following cDNA synthesis, samples were stored at −20°C. On the day of use, cDNA was diluted 1:10 in nuclease-free H_2_O, followed by transcript amplification according to manufactures instructions (Promega; GoTaq®, Cat. No A6001) using a ViiA 7 PCR system (Applied Biosystems) set to Standard cycling conditions. Gene expression data was normalized to *Actb*, and calculated using the 2^-ΔΔCT^ method. Primer pairs are listed in **Table 1**.

**Table 1.**
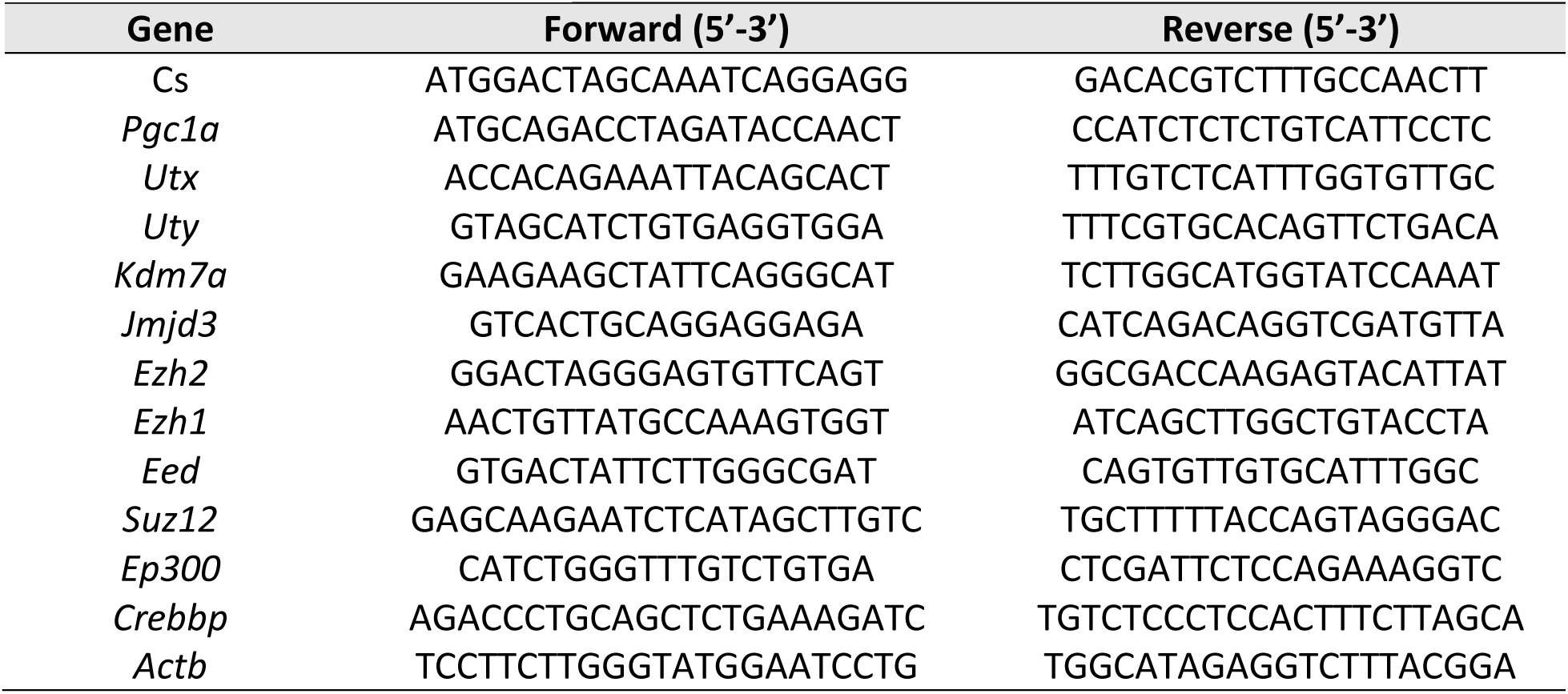
qPCR primer pairs.

### Statistics

Mean differences between two independent groups were analyzed with a Student’s two-tailed t-test, whereas one- and two-way repeated-measures ANOVAs followed by a Holm-Sidak post-hoc test were used where appropriate (GraphPad Prism 10.0.0). Statistical significance was determined at *P* < 0.05. All values are presented as mean ± S.E.M.

## Results

### Whole-Body Exercise Response

Average weekly running of male mice was similar to what we have previously found for C57BL/6J mice (**Fig. 1B**).^19^ Exercise volume for females was nearly 2-3X greater than that of male mice, in-line with previously documented sex-differences in mouse exercise capacity.^20^ Interestingly, the body weight of exercised males was reduced following running-wheel access relative to sedentary controls, whereas this was not the case for females (**Fig. 1C**). Of note, the body mass drop observed at the beginning of running wheel access was likely the result of stress encountered when moving from group- to single-housing conditions. Voluntary running had only a modest effect on body composition as measured by quantitative magnetic resonance (**Fig. 1D**). While not statistically significant, fat mass tended to be lower, with lean mass correspondingly increased in exercised mice of each sex. Consistent in direction with their changes in weekly body mass and composition following running, wet weights of retroperitoneal and inguinal adipose depots tended to be lower for exercised males, whereas no differences between sedentary and exercised females were seen (**Fig. 1E**). Voluntary running significantly increased soleus mass in females; similar results were seen for exercised males, although this was not statistically significant (**Fig. 1F**). No impact of exercise was seen for extensor digitorum longus mass in either sex.

**Figure 1.**
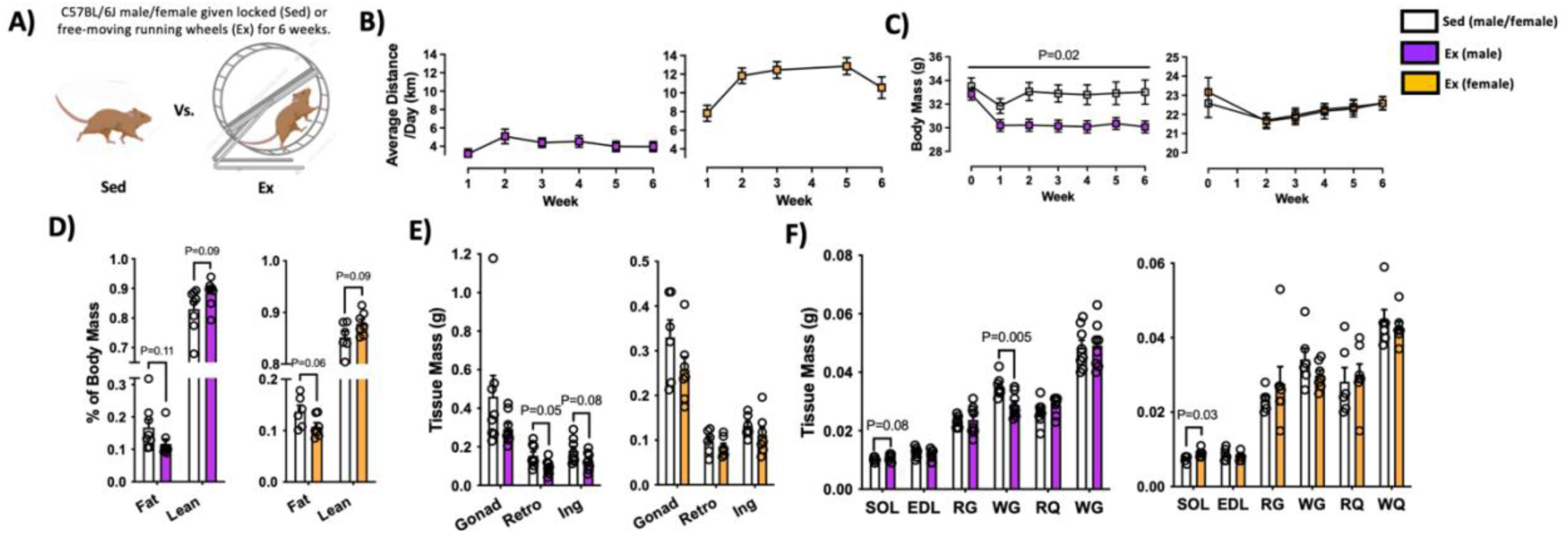
Body mass and composition response to 6 weeks of voluntary wheel running. **A)** Mice were given *ad libitum* access to free moving (Exercise; Ex) or locked (Sedentary; Sed) running wheels for 6 weeks. **B)** Average daily distance run (km) over six weeks. **C)** Weekly body mass (g) of Sed and Ex mice. **D)** Body composition (% fat/lean mass) as measured by quantitative magnetic resonance. **E)** Visceral and subcutaneous adipose depot mass (g). Gonad: gonadal; Retro: retroperitoneal; Ing: inguinal. **F**) Skeletal muscle mass (g). SOL: soleus; EDL: extensor digitorum longus; RG: red gastrocnemius; WG: white gastrocnemius; RQ: red quadricep; WQ: white quadricep. Values are mean ± S.E.M. n = 6-8/group.

### Gene Expression of H3K27-Modifying Enzymes

Exercise training has previously been shown to alter expression of several histone-modifying enzymes within skeletal muscle.^15,17^ Given the robust impact exercise has on muscle gene programs and the role H3K27 modifications play in mediating such response, we sought to determine if expression of H3K27 regulatory enzymes *themselves* were modified by physical activity. We chose to analyze an array of skeletal muscles differing in their oxidative/glycolytic capacities and fiber-type proportions. Of the representative slow/oxidative muscles analyzed (**Fig. 2 A-C**), expression of representative genes was most altered following 6 weeks of wheel running in male soleus. Gene expression of the mitochondrial master regulatory *Pgc1a* was significantly increased in exercised males only. Additionally, expression of the functionally homologous H3K27 acetyltransferases *Ep300* and *Crebbp* were both significantly upregulated in male SOL, whereas no other H3K27-modifying enzymes were changed in this tissue. *Eed*, a core component of the transcriptionally repressive polycomb repressive complex (PRC)-2, tended to be lower in the soleus of exercised females, while no other changes were seen in this muscle. No changes in the expression of marker genes were seen for this tissue or other slow/oxidative muscles analyzed.

**Figure 2.**
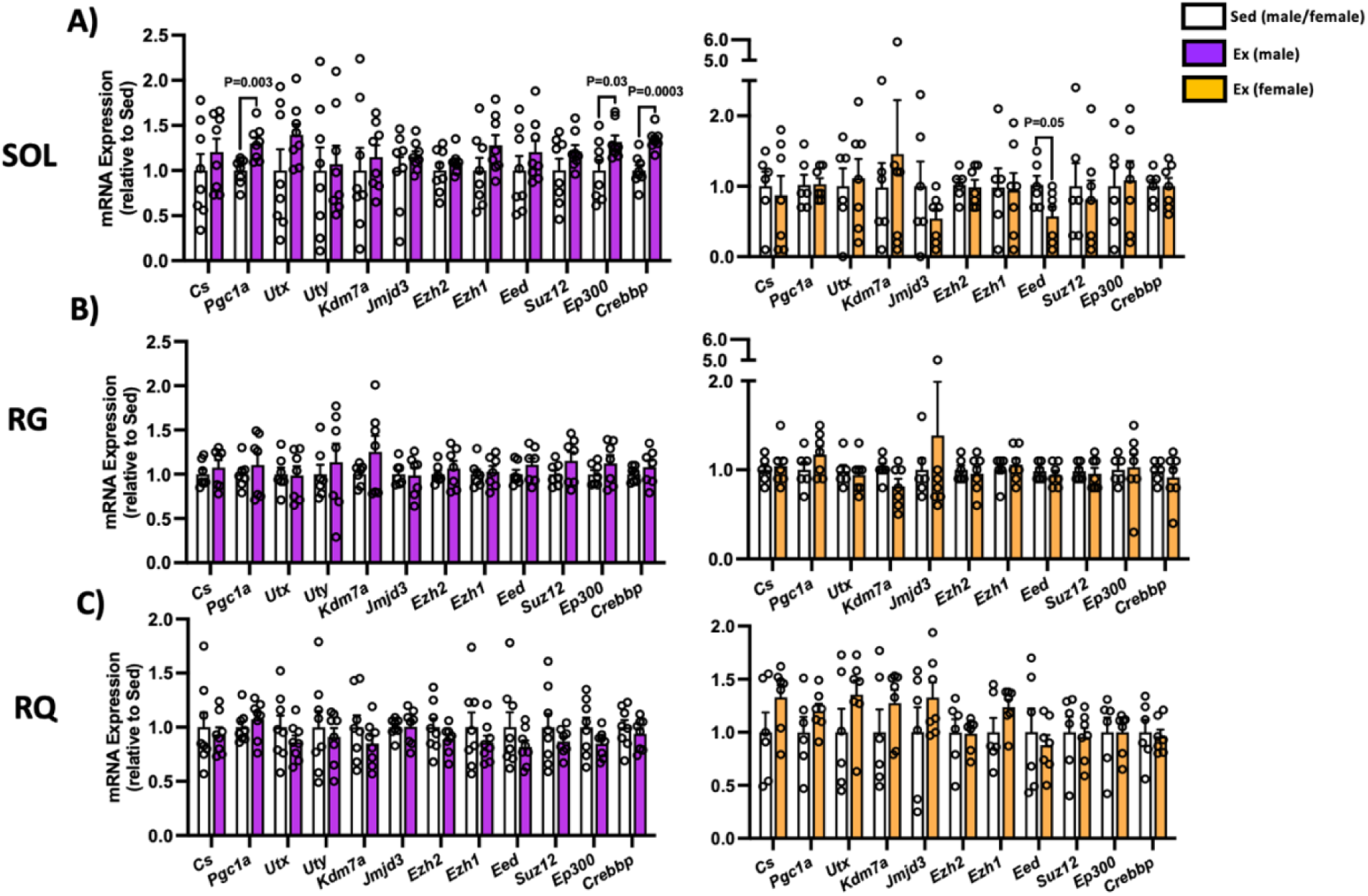
Gene expression response to 6 weeks of voluntary wheel running in representative slow/oxidative skeletal muscles. mRNA levels (expressed relative to Sed controls) of oxidative markers and regulators of H3K27 acetylation/methylation in **A)** soleus (SOL), **B)** red gastrocnemius (RG), and **C)** red quadricep (RQ). Values are mean ± S.E.M. n = 6-8/group.

Expression of representative genes remained relatively impervious to training in white gastrocnemius, although *Pgc1a* trended higher in exercised males (**Fig. 3 A**). Mitochondrial genes tended to be increased following 6 weeks of wheel running in male, but not female, white gastrocnemius. Several PRC-2 genes were elevated in male WG including the H3K27 methyltransferase *Ezh2* (although trending) and the core component *Suz12* (**Fig. 3 B**). Exercise only altered the expression of the H3K27 demethylase *Jmjd3* in WG of female mice.

**Figure 3.**
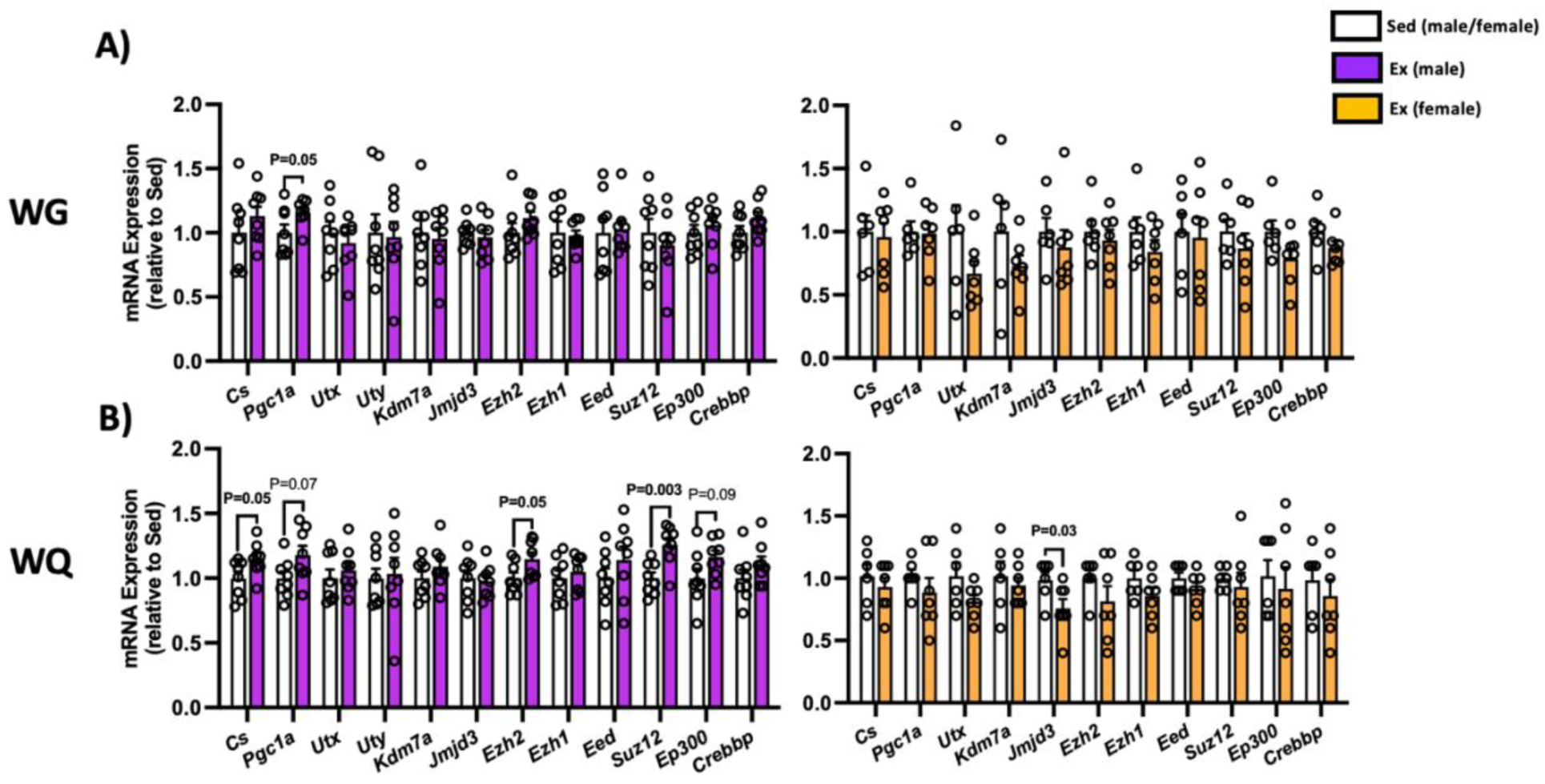
Gene expression response to 6 weeks of voluntary wheel running in representative fast/glycolytic skeletal muscles. mRNA levels (expressed relative to Sed controls) of oxidative markers and regulators of H3K27 acetylation/methylation in **A)** white gastrocnemius (WG), and **B)** white quadricep (WQ). Values are mean ± S.E.M. n = 6-8/group.

## Discussion

Skeletal muscle is uniquely capable of remodelling its structure and various contractile/metabolic properties in order to adapt to the demands placed upon it. Such plasticity is a function of complex signaling networks that decode context-specific information about physical activity into an appropriate response, which involves activation and suppression of adaptive gene programs. Modulating gene expression requires, among other factors, altering chromatin structure and thereby access of transcriptional machinery to gene regulatory regions (i.e. promotors, enhancers). Post-translational modifications to H3K27 are well known to regulate chromatin dynamics and transcriptional activation/repression. Several studies have shown that histone post-translational modifications and/or their regulatory enzymes are altered by exercise,^8,14–17^ suggesting an integral role in coordinating muscle adaptative remodelling. Our objective here was to examine whether the expression of major H3K27-modifying enzymes were responsive to physical activity in rodent muscles of varying metabolic and contractile characteristics. We found that 6 weeks of voluntary wheel running induced modest changes in gene expression of H3K27 acetyltransferases and PRC-2 core components in a sex- and muscle-specific manner.

Voluntary wheel running is a useful paradigm of exercise training that minimizes animal stress and enables mice to run during their natural circadian pattern.^20^ Weekly running volume of male mice was comparable to our previous work^19^ and others,^21–23^ although some have found higher volumes for C57BL/6J mice.^24–26^ In line with others,^27^ we found females to be superior runners relative to males throughout the training period. Unlike the males however, female body mass was surprisingly unchanged despite running nearly 2X more than males.

Importantly, we did not measure food intake in this study, which may have increased in females to compensate for the energetic demands of running. Regardless, modest reductions in fat mass and concomitant increase in lean mass were observed in runners of each sex. Although reduction in body fat percentage of male runners was consistent with the decline of several adipose depot wet weights, this was not the case for female runners, suggesting that other adipose depots were impacted by wheel running. The proportional increase in lean mass of running mice, regardless of sex, was consistent with the larger soleus wet weights, which has previously been shown in C57BL/6J mice following 7-weeks of weighted wheel running.^28^

Endurance exercise training is well known to enhance skeletal muscle oxidative metabolism and increase mitochondrial biogenesis. In agreement with this, we found oxidative markers (e.g. *Cs, Pgc1a*) to be increased in skeletal muscles of males only; this was initially surprising given the markedly greater running volume of females. However, a recent meta-analysis found biochemical adaptations to exercise training are less pronounced in female mice.^29^ Initial knockout studies of *Pgc1a* revealed defects in muscle mitochondrial programming,^30^ whereas physiological overexpression in skeletal muscle mimicked many biochemical, molecular, and functional adaptions of endurance training.^31,32^ Despite this, several studies revealed *Pgc1a* is not obligatory for exercise-induced mitochondrial biogenesis,^33,34^ likely owing to functional redundancy in pathways controlling oxidative metabolism. Furthermore, while expression of oxidative genes/proteins are common markers of endurance training efficacy, they are by no means the only relevant adaptive readouts given the pleiotropic effect of exercise. Thus, it is unlikely that no functional adaptation(s) occurred in the skeletal muscles of female runners.

Mounting evidence indicates post-translational modifications of histones regulate homeostatic and adaptive programming of skeletal muscle. Deacetylation of histone lysine residues is well-known to facilitate transcriptional repression, and inhibiting various histone deacetylases (HDACs) in rodents can broadly recapitulate exercise adaptions, including enhancing mitochondrial biogenesis and promoting fast-to-slow fiber type transition.^35–38^ Importantly, exercise causes the nuclear export of HDACs in muscle,^10,39^ suggesting transcriptional activation of exercise-responsive genes is aided by HDAC subcellular redistribution. Although histone acetylation and subsequent expression of metabolic genes are enhanced by HDAC inhibition,^37^ it remains unclear how much of this effect is driven by H3K27ac *per se*, particularly given that no H3K27-specific deacetylase has been identified. Adaptive muscle remodelling via HDAC inhibition could result from greater occupancy/access to H3K27 by histone acetyltransferases. Acetylation of H3K27 is entirely dependent on the functionally homologous proteins p300 (*Ep300*) and cAMP-response element binding protein (CREB) binding protein (CBP; *Crebbp*).^40–42^ Importantly, prolonged endurance training of both humans^14^ and rodents^8^ markedly alters genome-wide distribution of H3K27ac, suggesting p300/CBP activity is recruited to co-ordinate adaptive gene programing in response to exercise. In line with this, we provide evidence here that the capacity of p300/CBP may be enhanced in the trained state (**Fig. 2 A**). Given their considerable redundancy, tissue-specific single knockouts for p300 or CBP maintain muscle homeostasis and metabolic responses to prolonged wheel running,^43,44^ whereas inducible double knockout profoundly impairs muscle function resulting in death.^45^ As such, the exercise-responsive gene programs regulated by p300/CBP and their necessity in mediating remodelling of muscle remain unclear.

Contrary to acetylation, methylation of H3K27 results in chromatin compaction and gene silencing. H3K27me1-3 formation is a function of the polycomb repressive complex (PRC)-2, which is comprised of the core components embryonic ectoderm development (*Eed*), suppressor of zeste (*Suz12*), and the catalytic methyltransferase enhancer of zeste homolog 2 (*Ezh2*). Gene expression of several PRC-2 components were elevated in fast/glycolytic muscle by wheel running (**Fig. 3 B**). Several studies have shown global H3K27me3 accumulates in muscle following acute and prolonged exercise training in both human^46^ and rodent skeletal muscle.^11,16^ While seemingly contrary to its canonical role in transcriptional repression, H3K27me3 is required for inducing molecular adaptations with exercise, as pharmacological inhibition of PRC-2 activity impairs endurance adaptations in rodent muscle.^11^ Importantly, such inhibition may prevent formation and maintenance of bivalent (poised) chromatin regions, which are marked by both transcriptionally repressive (i.e. H3K27me3) and activating (i.e. H3K4me3) marks. Moreover, promoting H3K4 methylation within rodent muscle mimics endurance training, including enhancing slow-twitch programming and improved running performance.^17,47^ Here, we show here that wheel running enhanced gene expression of the methyltransferase *Ezh2* along with *Suz12*, which is required for maintaining PRC-2 integrity and enzymatic function.^48^ While mechanistically unclear from the present study, increasing PRC-2 capacity may facilitate transcriptional repression of some genes while simultaneously maintaining poised regions vital for facilitating endurance adaptations.

While we show modest gene expression changes in several regulators of H3K27 PTMs, many remained unaffected following 6 weeks of wheel running, particularly within female skeletal muscles; several reasons may account for this. Although voluntary wheel running reduces animal stress and enables long term exercise studies without a need for continuous monitoring, there is an obvious trade-off in lack of control for exercise frequency, intensity, duration, and circadian timing of running. Secondly, although we collected tissues post-training where we were likely to observe the greatest increase in classic mitochondrial markers with exercise, we cannot rule out that changes in H3K27 modifying genes occurred during earlier stages of the experiment. Furthermore, we cannot rule out that changes in genomic location, enzyme activity or their subcellular distribution were not affected by wheel running, independent of gene expression. Certainly, several studies have shown endurance training to modulate loci-specific and/or global changes in H3K27ac/me3 enrichment. ^8,14–16^ Ultimately, muscle-specific loss-of-function studies are required to delineate which gene programs are regulated by individual H3K27-modifiers with exercise training.

Exercise training improves overall health and muscle functioning by remodelling various structural and metabolic programs. Such remodelling requires altering the chromatin landscape to support activation and suppression of adaptive gene programs. Chromatin dynamics are regulated by numerous factors like post-translational modifications to various histone residues, including H3K27. Here, we provide the first sex-specific investigation into the impact of prolonged exercise on expression of enzymes regulating the H3K27 transcriptional switch within skeletal muscle. We found that 6 weeks of voluntary wheel running altered gene expression of epigenetic machinery responsible for regulating H3K27ac/me3 enrichment in both a muscle- and sex-specific manner. Our findings add to a growing body of evidence that histone modifications serve as a regulatory node co-ordinating muscle plasticity.

